# A New Halogenated Solvent For Ex Vivo Magnetic Resonance Imaging

**DOI:** 10.1101/2024.06.05.597589

**Authors:** Nicolas Honnorat, Mariam Mojtabai, Jinqi Li, Morgan Smith, Sudha Seshadri, Kevin Bieniek, Mohamad Habes

**Affiliations:** Glenn Biggs Institute for Alzheimer’s and Neurodegenerative Diseases, University of Texas Health Science Center at San Antonio, San Antonio, Texas, USA; Research Imaging Institute, University of Texas Health Science Center at San Antonio, San Antonio, Texas, USA; Department of Pathology, University of Texas Health Science Center at San Antonio, San Antonio, Texas, USA

## Abstract

Magnetic Resonance Imaging (MRI) is a prominent non-invasive imaging technique. Structural MRI, the most common MRI modality, interacts with the hydrogen nuclei in samples, also called protons. When using structural MRI to scan ex vivo tissues, the biological samples are often placed in proton-free liquids containing no hydrogen atoms to obtain clean images that do not require background removal. Several proton-free liquids have been used during the last two decades, but they are all per- and polyfluoroalkyl substances (PFAS), and PFAS have recently been recognized as a significant environmental concern. To find replacement solutions, new families of proton-free liquids without fluorine atoms will need to be investigated for stability and safety and to check potential effects on subsequent tissue staining. The present work is a preliminary step in that direction. We validate the MRI properties of a broadly used, affordable, non-PFAS, proton-free solvent that has, as far as we know, never been considered for ex vivo MRI scanning.

## 1 Introduction

Magnetic Resonance Imaging (MRI) is a prominent non-invasive imaging technique that uses strong magnetic fields and radiofrequency (RF) waves to interact with the magnetic spin of molecules and atoms [1]. 3D visualizations are created by analyzing the signals emitted by these spins as they realign with a reference magnetic field after being disturbed by RF pulses [1]. Structural MRI, the most common MRI modality, refers to specific pulse sequences interacting with the hydrogen nuclei in samples, particularly abundant in water and fat in biological tissues [1]. These hydrogen nuclei are also called *protons*. When using structural MRI to scan ex vivo tissues, the biological samples are often placed in *proton-free* liquids containing no hydrogen atoms to obtain clean images that do not require background removal [2, 3, 4, 5]. Perfluoropolyethers (PFPE) have been used for this task for more than twenty years [2]. The fluids sold under the commercial names Fomblin and Galden by the Solvay company appear the most often in the literature [2, 6, 3], but their competitor Krytox sold by the DuPont/Chemours company has been used as well [4].

Since perfluoropolyethers such as Fomblin can be very expensive [3], other perfluorinated compounds have been recently proposed: the fluorocarbons sold under the FluorInert brand name by the 3M company, and more specifically, the FluorInert FC-77 [7, 8, 9] and the FluorInert FC-3283 [3]. Unfortunately, most of these fluids often used to cool electronics systems have a large atmospheric lifetime associated with global warming potentials several thousands of times larger than carbon dioxide [10, 11]. New fluorocarbons were proposed to fix the issue, such as the fluoroketone sold under the brand names Novec 1230/649 by 3M and the product identifier FK-5-1-12 by competitors. These fluids possess physical properties similar to FluorInert fluorocarbons but moderate global warming potentials, and they have just been successfully tested in MRI scanners [5]. But an issue has emerged in the last couple of years. Perfluoropolyethers (PFPE), FluorInert fluorocarbons, and fluoroketones are all per- and polyfluoroalkyl substances (PFAS), and PFAS have recently been recognized as a significant environmental concern [12]. Their stability make them very persistent [13], and environmental protection agencies are currently trying to restrict their production and their use by determining which non-essential PFAS uses should be terminated [14]. As a result, all Novec and Fluorinert products will be discontinued by their manufacturer by the end of 2025 [15].

This last development will have a strong impact in the scientific field, as most of the proton-free liquids used so far are under the threat of being abruptly discontinued in the coming years [3]. To find replacement solutions, new families of proton-free liquids without fluorine atoms will need to be investigated for stability and safety and to check potential effects on subsequent tissue staining [3]. The present work is a preliminary step in that direction. We validate the MRI properties of a broadly used, affordable, non-PFAS, proton-free solvent that has, as far as we know, never been considered for ex vivo MRI scanning.

## 2 Method

### 2.1 The New Halogenated Solvent

Tetrachloroethylene, also known as perchloroethylene and PCE, is a common halogenated solvent used for dry cleaning, textile processing, and metal degreasing [16, 17]. It is a transparent liquid heavier than water (density 1.622 grams per cubic centimeter, molar mass 165.82 g/mol), with a melting point around −22 ^*o*^C and a boiling point at 121.1 ^*o*^C.

Tetrachloroethylene presents several promising characteristics for ex vivo magnetic resonance imaging. First, the tetrachloroethylene solubility in water is low: close to 0.2 g/L at room temperature [18]. Then, tetrachloroethylene has a moderate vapor pressure, of 0.020338 atm or 2060.7 Pa at 295 Kelvins (21.85 ^*o*^C) [18], smaller than the vapor pressure of water at the same temperature: 2620.8 Pa [19]. In addition, the molar magnetic susceptibility of tetrachloroethylene 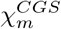(*PCE*) was estimated at −81.6×10^−6^ cm^3^/mol which corresponds to a dimensionless volume magnetic susceptibility 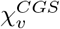(*PCE*) of −7.983 ×10^−7^ [20, 21], and this value is close to the volume magnetic susceptibility of water and biological samples: 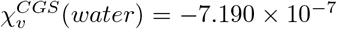 [22]. As a result, tetrachloroethylene is unlikely to evaporate quickly, to diffuse swiftly in biological samples, or create strong magnetic resonance imaging artifacts.

### 2.2 Sample Preparation

Two similar glass bottles were used. In the first bottle, one hundred milliliters of a 10% buffered formalin solution were prepared [23], while the second bottle was filled with one hundred milliliters of tetrachloroethylene (Sigma-Aldrich).

A piece of beef meat was added inside each bottle, the bottles were closed, and placed in separate plastic bags that were zipped to prevent potential vapor leaks. This preparation is illustrate in Figure 1.

**Figure 1:**
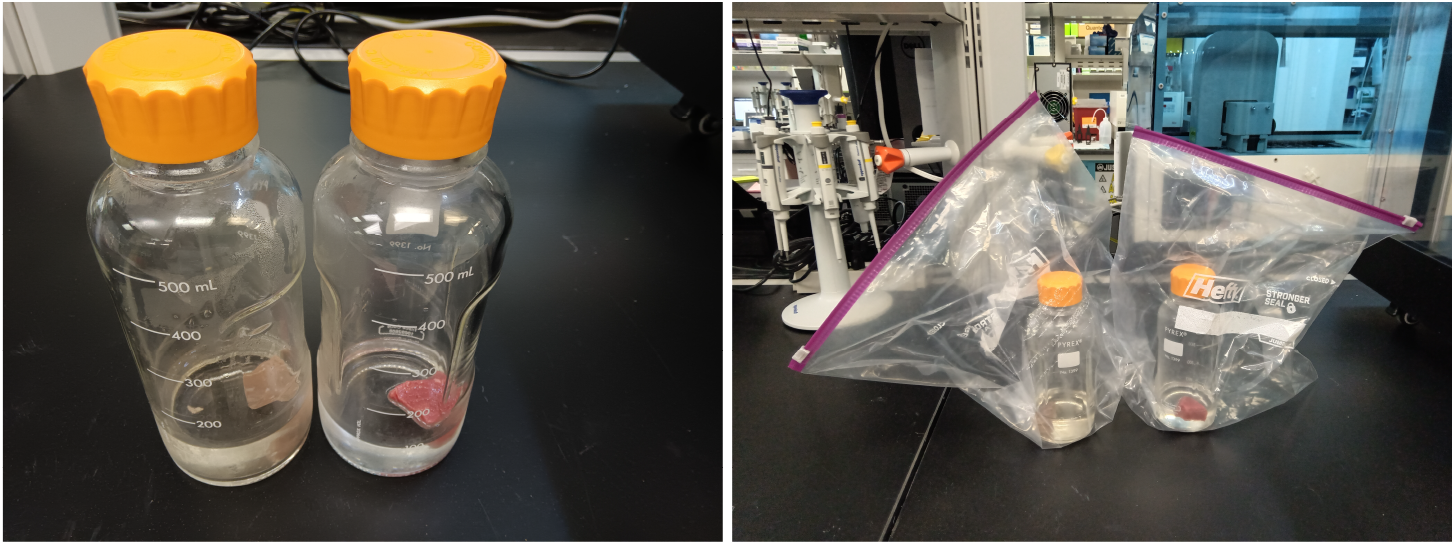
Experimental setting: meat samples were placed in two different glass bottles: one filled with formalin (on the left) and one filled with tetrachloroethylene. The bottles were placed in zipped plastic bags to prevent vapor leaks. These pictures were taken two days after the scan and the meat sample in the formalin bottle has oxidized and fixed while the sample in the tetrachloroethylene bottle remained in its original state.

### 2.3 MRI Scans

A Siemens TIM TRIO 3T whole-body MRI scanner was used to produce images of the two bottles at the same time. Two short scans were acquired: a T1-weighted MRI scan was acquired at a spatial resolution of 1.875mm × 1.875mm × 0.54mm, with a repetition time of 1450 ms, an echo time of 3.14 ms, an inversion time of 766 ms, a flip angle of 13^*o*^, and for a single average (no averaging). Then, a T2-weighted MRI scan was acquired at a spatial resolution of 1.563mm × 1.563mm × 0.8mm, with a repetition time of 4000 ms, an echo time of 23 ms, a flip angle of 120^*o*^, and without averaging.

## 3 Results

### MRI Scans

As shown in Figure 2, the tetrachlorethylene was completely transparent in both T1-weighted and T2-weighted scans.

**Figure 2:**
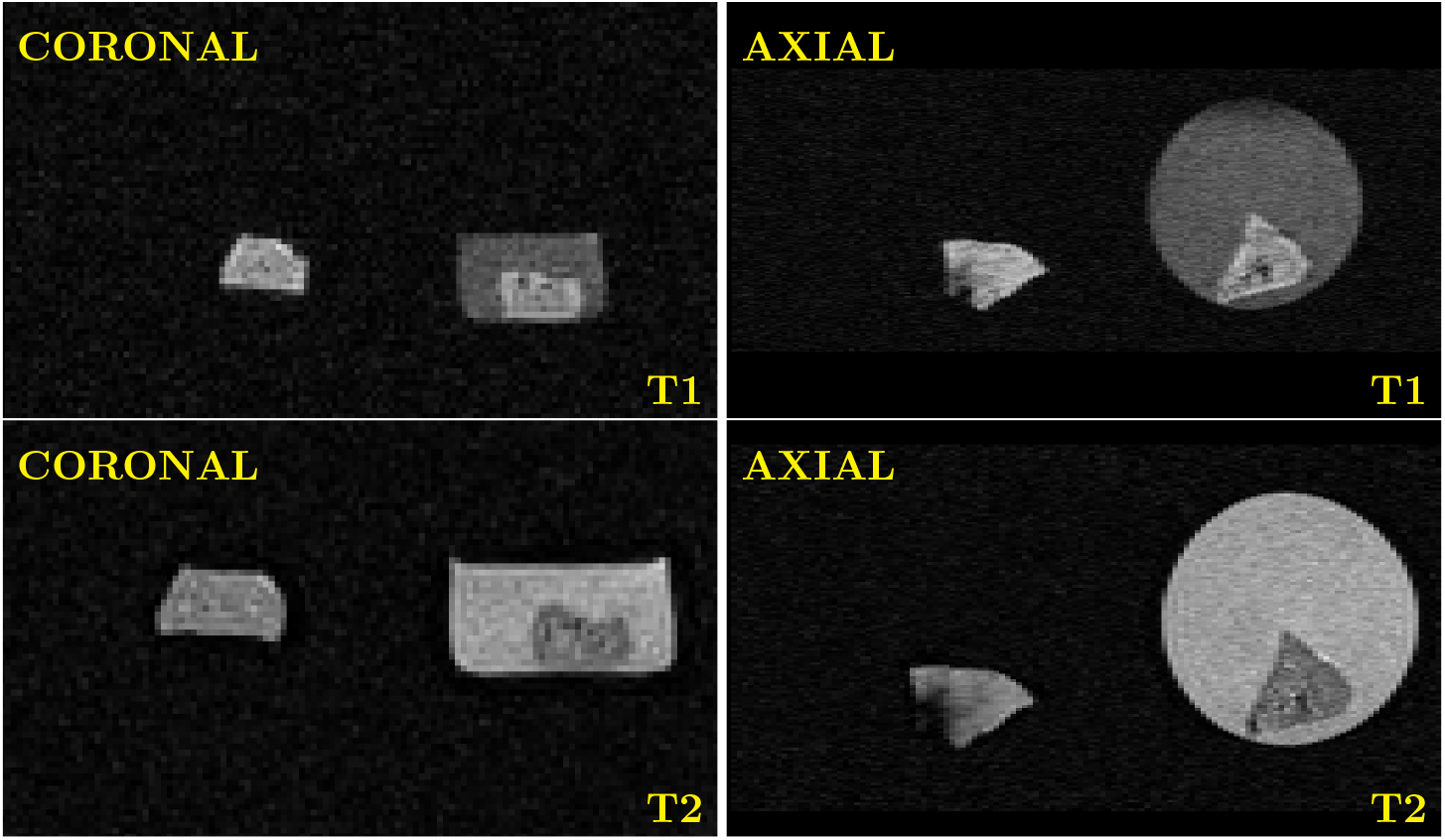
Coronal and axial views acquired by the T1-weighted and the T2-weighted scans. The two pieces of meat produced similar MRI intensity, but the formalin is the only visible fluid. The glass bottles, the plastic bags, and the air are also invisible.

### Samples Evolution

After eight days, there was no apparent change or decay of the meat sample inside the bottle filled with tetrachloroethylene. On the opposite, the sample in the formalin bottle has oxidized and fixed. This effect is already visible in the pictures taken two days after the scan reported in Figure 1. After eight days, there was no apparent change of fluid level in the bottles and no noticeable vapor leaks. These observations suggest that the tetrachloroethylene has not seriously damaged the meat sample and not strongly evaporated.

## 4 Conclusion

Tetrachloroethylene is a promising proton-free liquid for ex vivo imaging. More investigations are required to check if the fluid alters the histological properties of biological tissues. But these experiments will need to be carried out for samples already fixed in formalin instead of fresh meat.

